# Mapping the landscape of anti-phage defense mechanisms in the *E. coli* pangenome

**DOI:** 10.1101/2022.05.12.491691

**Authors:** Christopher Vassallo, Christopher Doering, Megan L. Littlehale, Gabriella Teodoro, Michael T. Laub

## Abstract

The ancient, ongoing coevolutionary battle between bacteria and their viral predators, bacteriophages, has given rise to sophisticated immune systems including restriction-modification and CRISPR-Cas. Dozens of additional anti-phage systems have been identified based on their co-location within so-called defense islands, but whether these computational screens are exhaustive is unclear. Here, we developed an experimental selection scheme agnostic to genomic context to identify defense systems encoded in 71 diverse *E. coli* strains. Our results unveil 21 new and conserved defense systems, none of which were previously detected as enriched in defense islands. Additionally, our work indicates that intact prophages and mobile genetic elements are primary reservoirs and distributors of defense systems in *E. coli*, with defense systems typically carried in specific locations, or hotspots. These hotspots in homologous prophages and mobile genetic elements encode dozens of additional, as-yet uncharacterized defense system candidates. Collectively, our findings reveal an extended landscape of antiviral immunity in *E. coli* and provide a generalizable approach for mapping defense systems in other species.

## Introduction

Bacteriophages (or simply, phages) are an extraordinarily diverse and ubiquitous class of viruses that pose a nearly constant threat to bacteria. Phages are the most abundant biological entity on the planet, with estimates of 10^31^ particles that drive the daily turnover of ∼20% of all bacteria in some environments^1,2^. Bacteria and their viral predators are locked in a perpetual coevolutionary battle, leading to the emergence of sophisticated mechanisms by which phage manipulate and exploit their hosts and an equally diverse set of bacterial immune mechanisms collectively referred to as anti-phage defense systems^3^. These immunity systems include both innate mechanisms such as restriction-modification systems and adaptive mechanisms such as CRISPR-Cas. Recent studies have begun to identify many new defense systems, but the full inventory likely extends well beyond what is currently defined.

Identifying additional anti-phage defense systems promises to provide new insight into the ancient coevolutionary conflict between viruses and their hosts. Recent work has found that many defense systems have homologs with similar function in eukaryotic innate immunity, indicating a potentially ancient origin and cross-kingdom conservation of many immune systems^4–6^. Additionally, prior studies of anti-phage defense have produced precision molecular tools such as CRISPR and restriction enzymes, so the discovery of new immune mechanisms may enable new tools for manipulating cells and genomes. Finally, there is growing interest in using phages to treat antibiotic-resistant bacterial infections and to manipulate microbiomes^7–9^. A more complete understanding of the diverse mechanisms by which bacteria defend themselves may be critical for these endeavors^10^.

Multiple groups have previously used computational methods to identify uncharacterized defense systems, building off of the observation that anti-phage defense systems often cluster in bacterial genomes in high density, forming so-called “defense islands”^11–13^. However, not all defense systems may be detectable in these guilt-by-association approaches. For systems that are rare or not widely conserved it may be difficult to detect enrichment within defense islands, and not all defense systems necessarily associate with defense islands. Additionally, candidates identified computationally must be expressed in a model laboratory organism and then tested against a panel of phages. Some systems may not work in a heterologous host or protect against the phages examined, and demonstrating that a given system provides defense in its native context is typically not tested or even possible. We reasoned that an experimental selection scheme to uncover antiviral proteins may reveal new insights into bacterial immunity, including identifying defense systems that remain uncharacterized and the relative frequency of the different genomic contexts of these bacterial immune systems (*i*.*e*. defense islands, mobile genetic elements).

To this end, we took a functional metagenomic approach to map the range of defense systems in the *E. coli* pangenome. We built a fosmid library from a diverse set of 71 wild isolate *E. coli* strains and then introduced this library into an *E. coli* K12 derivative. We then challenged the library with three diverse lytic phages, T4, T7, and λ_vir_, and directly selected for phage resistance. We then used a high-throughput, deep sequencing-based approach to pinpoint 21 novel anti-phage defense systems within the fosmids (Fig. 1). Although some of these systems are associated with defense islands, most are not. Instead, our work supports a major role for resident prophages and mobile genetic elements as primary contributors to bacterial immunity. Some of the systems identified feature proteins or domains rarely or never previously implicated in phage defense. Similarly, we find and characterize several atypical toxin-antitoxin (TA) systems that provide potent defense, underscoring the likely widespread role that these systems play in bacterial immunity. Importantly, in addition to revealing the landscape of phage defense in *E. coli*, our work also provides a robust screening methodology that can now be adapted to systematically identify phage defense mechanisms in virtually any bacterial genome or metagenomic sample.

**Figure 1.**
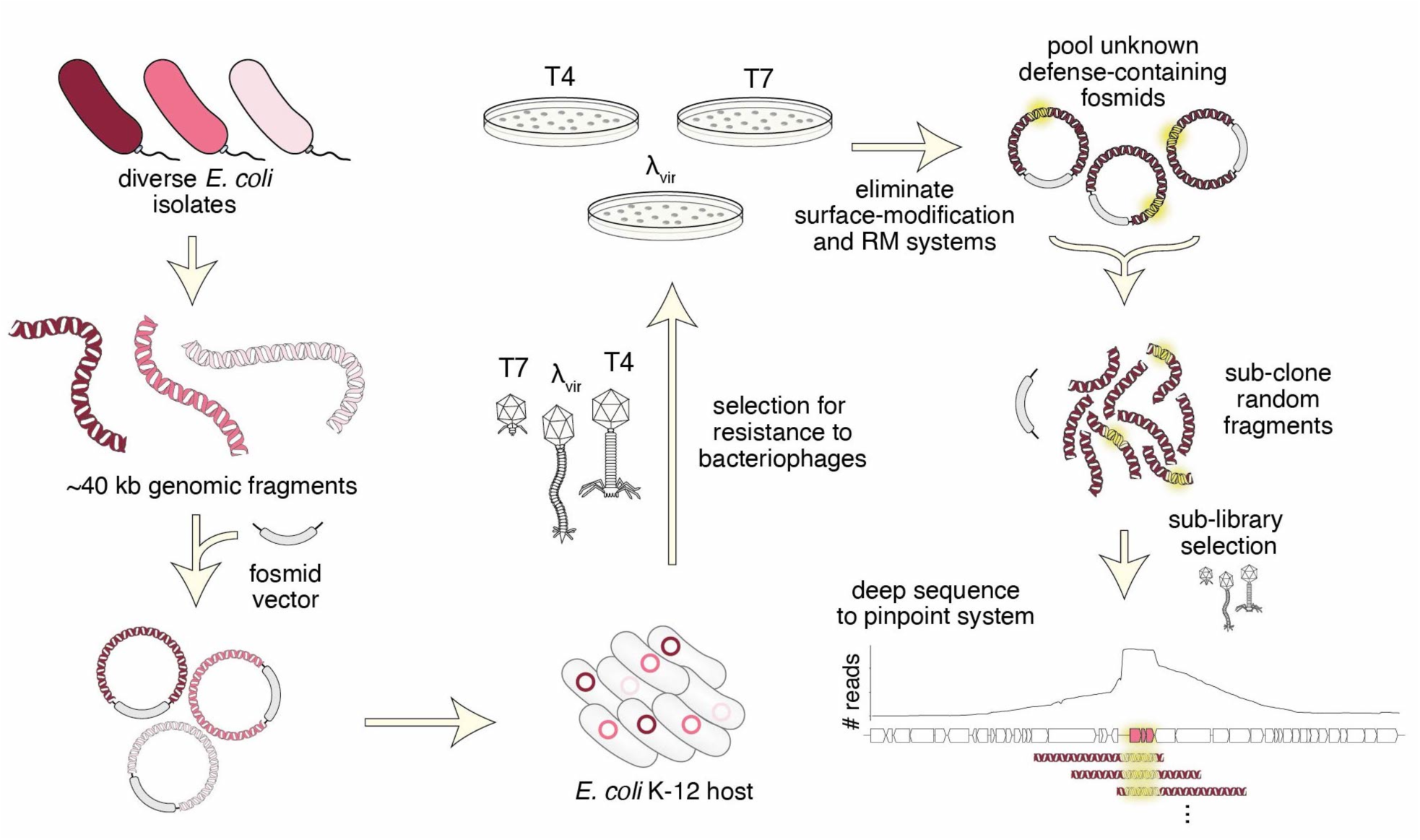
Selection strategy for identifying novel phage defense systems. A fosmid library of random ∼40 kb fragments of genomic DNA from 71 *E. coli* strains was transformed into an *E. coli* K-12 host and then challenged with three different phages. Survivors were isolated and fragments mapped to their genome sequence. After eliminating duplicates, clones affecting adsorption, and clones harboring restriction-modification or known systems, the unique fosmids corresponding to each phage selection were used to construct plasmid libraries, which were subjected to a second selection. Surviving clones were deep-sequenced and candidate defense loci pinpointed by mapping sub-library reads to genome sequences of the original fosmid inserts.

## Results

### Identification of novel anti-phage defense systems

To sample the immune landscape of *Escherichia coli*, we collected a diverse set of wild isolate strains from the ECOR collection as well as 19 clinical isolates^14^ (71 strains in total), all with available draft genome sequences (Fig. S1). The ECOR collection is a set of strains curated to span the phylogenetic diversity of the species^15^. Together, the 71 strains collected encode 21,149 unique gene clusters, of which > 10,000 exist in only one or two strains (Fig. S1). From genomic DNA, we constructed a 100x-coverage library of fosmids, each harboring an ∼40 kb genome fragment, in EPI300, a derivative of *E. coli* K12. We used large-insert fosmids to minimize the size of the library, to include potentially large defense systems, and because the copy number is maintained around one, minimizing false positives due to overexpression.

Many anti-phage defense systems work by an abortive infection (Abi) mechanism in which an infected cell sacrifices its viability to prevent phage replication and thereby protect uninfected cells in a population^16^. Thus, it is impossible to directly select for clones containing an Abi-based defense system because the infected cell dies. Instead, we used a selection strategy, historically known as the *tab* (<u>T</u>4 <u>ab</u>ortive) method^17^. In short, cells harboring the gDNA library are mixed with phage in a structured medium (soft agar) at varying concentrations of phage (Fig. 2a). At intermediate phage concentrations, individual clones from the library can grow and form small populations before encountering a phage particle. Any such micro-colonies that harbor an Abi defense system can be infected, but the initially infected cell will die without producing progeny phage, enabling the rest of the population to survive and produce a colony. Thus, our screening approach allows the identification of both Abi and conventional defense systems.

**Figure 2.**
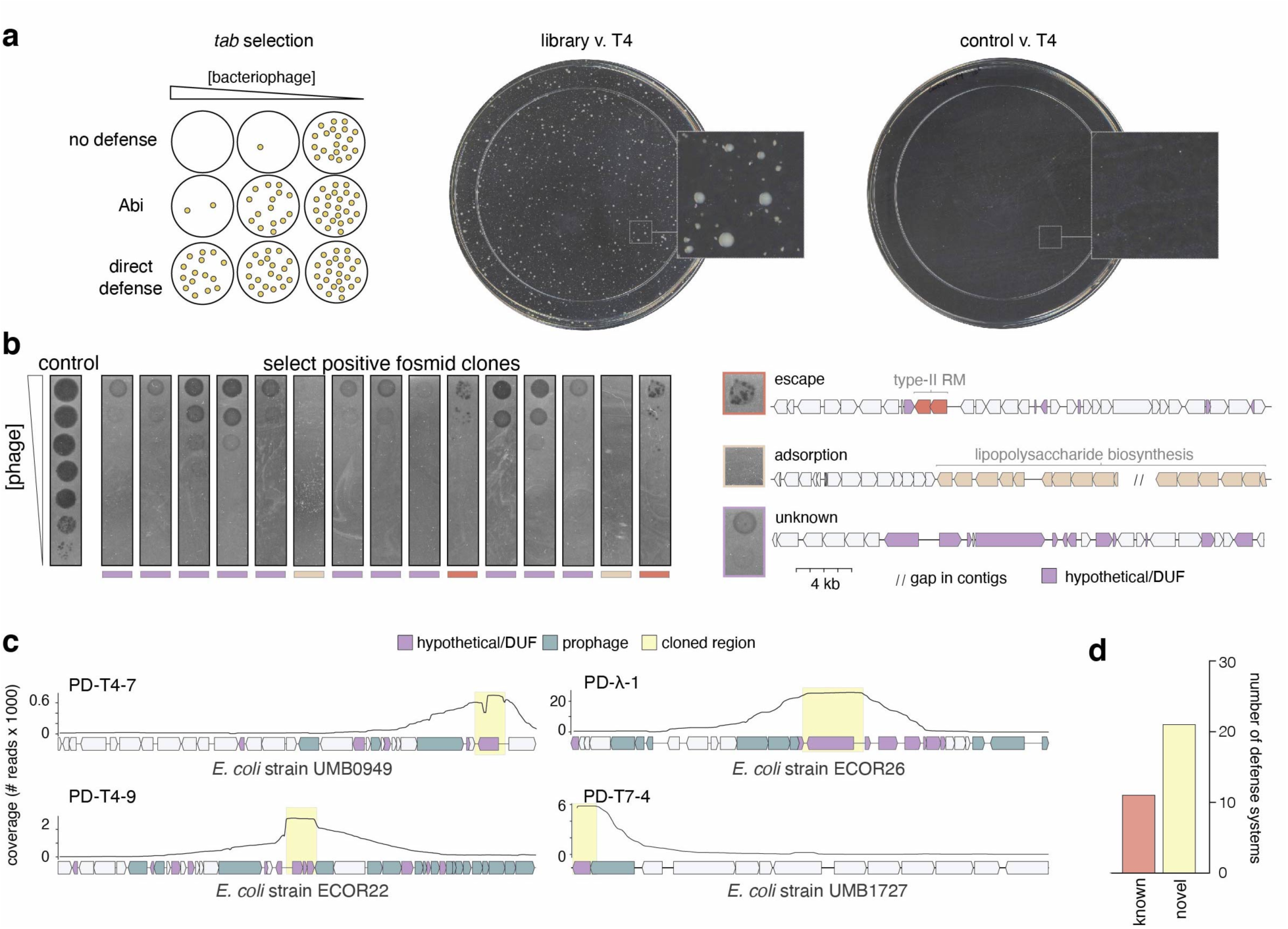
Identification of novel phage defense systems. (**a**) (*Left*) Schematic of the *tab* selection method. At intermediate concentrations of phage, *tab* selection facilitates the survival of cells with either abortive infection or direct defenses. (*Right*) Examples of T4 selection plates for cells containing the fosmid library or an empty vector control. (**b**) (*Top*) Ten-fold dilutions of λ_vir_ phages on lawns of a sample of 15 positive clones from the λ_vir_ screen. Multiple phenotypes were observed, including reduction of plaquing with individual escape plaques indicative of a restriction-modification system, no lysis at any concentration of phage typically reflecting a loss of adsorption, or reduction of plaquing, generally indicative of a phage defense system. Examples of fosmid inserts corresponding to exemplar phenotypes in (**b**) with relevant genes colored. (**c**) Examples of read coverage (100 bp moving average) from deep-sequencing of sub-libraries generated from positive fosmid clones with maxima delineating defense system candidates. Genes were colored or shaded as indicated at the top. (**d**) Summary counts of defense systems identified.

Using this general strategy, we challenged cells harboring the fosmid library with three lytic phages; T4, λ_vir_, and T7, each representing a major class of Caudovirales, the tailed bacteriophages (Fig. 1). From each selection, we isolated approximately 90 surviving colonies and then sequenced the ends of the vector insert in each clone to identify the genomic region and strain of origin of each fragment. For each positive clone we then measured the efficiency of plaquing (EOP), the ratio of plaques formed by a phage plated on the positive clone compared to a phage-sensitive control strain (Fig. 2b). In some cases, these EOP assays revealed nearly complete protection even at phage titers of >10^9^ PFU ml^-1^. Upon initial sequencing, representative clones with this phenotype consistently encoded lipopolysaccharide or capsule biosynthesis genes. This phenotype is consistent with a loss of adsorption that results from modification of the core surface properties of the cell (Fig. 2b, S1). Other positive clones produced protection accompanied by a high rate of escape plaques, a phenotype seen with many restriction-modification systems (Fig. 2b). Escape plaques can arise with many defense systems, but are well known to arise at high frequencies for RM systems due to epigenetic escape^18^. Upon sequencing, the clones exhibiting high frequency escape plaques indeed encoded RM systems. Thus, clones with these two phenotypes were excluded from further analyses (see Methods). From 257 initial clones, 117 and 9 were eliminated as likely resulting from changes in cell surface properties affecting adsorption or RM-mediated defense, respectively. After also accounting for redundancy in the remaining 131 clones, we had 43 clones that we hypothesized encode novel defense systems.

To pinpoint possible defense systems, we pooled the 3 sets of remaining fosmids corresponding to each phage and generated random 6-12 kb fragments. These fragments were then sub-cloned into a low-copy plasmid vector to create three high-coverage sub-libraries. Cells harboring these high-coverage sub-libraries were then selected for resistance to their respective phage, and plasmids from positive clones were sequenced by Nanopore long-read sequencing. The sub-library clones that survived selection each contained the originally selected defense system flanked by random lengths of adjacent DNA from the original fosmid insert. Thus, when reads were mapped back to the genome fragment in a given fosmid, the coverage maxima typically delineated the boundaries of each candidate defense system (Fig. 2c). In some cases, this identified previously-characterized systems, including type III and IV RM systems and an Old-family endonuclease, or non-defense genes that account for phage resistance, such as *malI*, a regulator of the lambda receptor LamB (Table S1). After discarding such cases, there were 21 unique candidate defense systems, with 10, 6, and 5 systems from the selections for T4, λ_vir_, and T7, respectively (Fig. 2d, Table S2). Each candidate system was provisionally named with a PD prefix, for phage defense, followed by the phage used for selection and a unique number.

### Validation of candidate defense systems

To validate these novel defense systems, we cloned each candidate ORF or operon into a low-copy vector under the control of its native promoter in wild-type MG1655 (Table S2). We confirmed that each system did not affect phage adsorption (Fig. S2b) and then challenged each with a panel of 10 diverse phages (Fig. 3a). Each candidate system was confirmed to substantially reduce the EOP for the phage originally used to select the system, and often others. In some cases, a given defense system did not change the EOP of a phage but instead produced smaller plaques. Although most defense systems were relatively specific, protecting against only a few phages, some systems provided relatively broad protection, such as PD-λ-5, which affected EOP or plaque size for all but one of the phages tested. The 10 systems selected to defend against T4 also generally protected against the other, related T-even phages, T2 and T6. Most systems protected most strongly against the phage originally used to select it, but with some exceptions. For instance, PD-λ-5, PD-T7-1, and PD-T7-3 protected more strongly against the T-even phages than against the λ_vir_ and T7 phages used to identify them. The fact that these systems provided robust defense against T4, but were not identified in the T4 selection, indicates that our screen was not saturating and that the systems identified represent only a subset of the defense systems in the original 71 *E. coli* isolates used.

**Figure 3.**
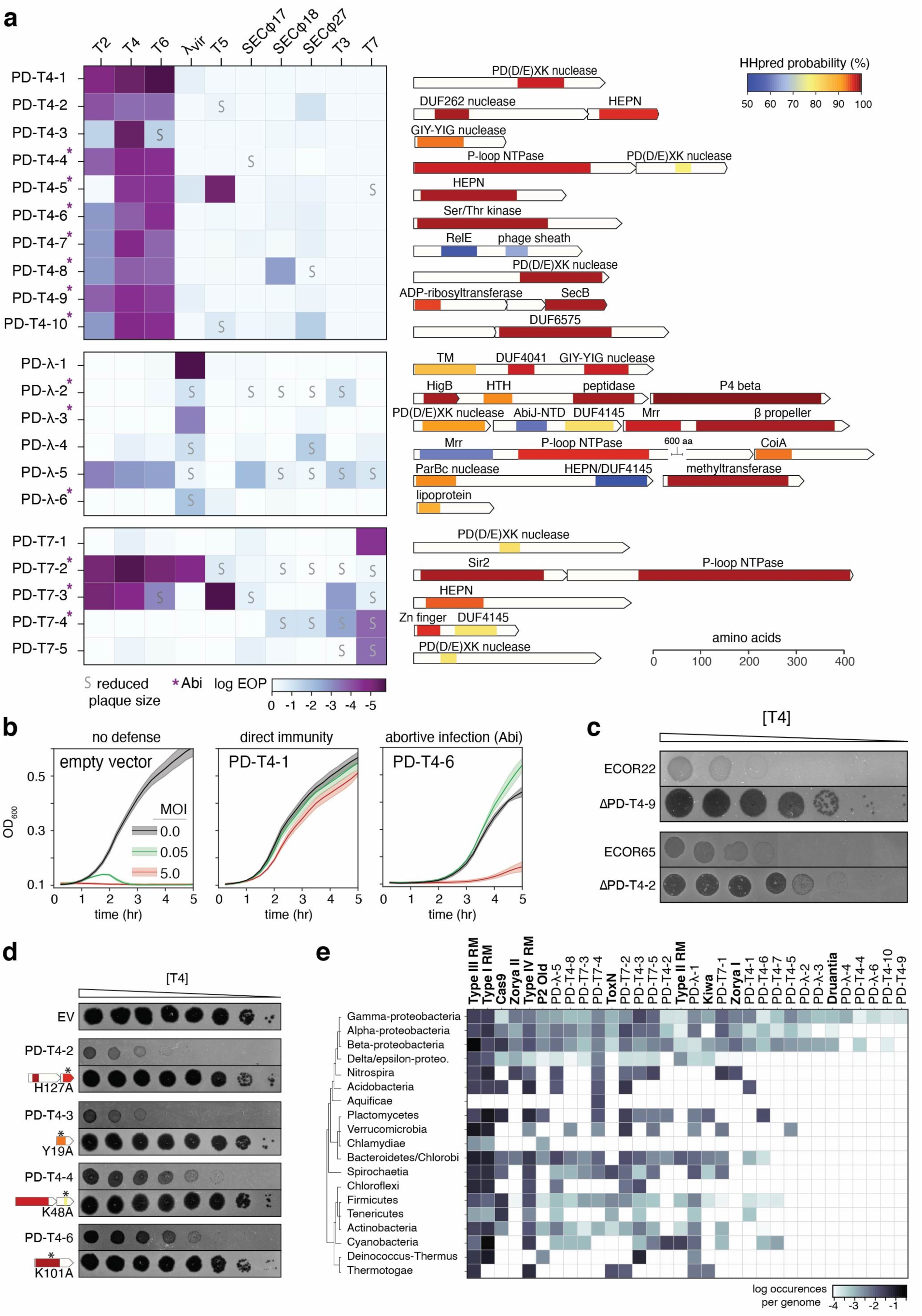
Summary and annotation of 21 novel defense system loci. (**a**) Each defense system was cloned into a low-copy plasmid with its native promoter and the efficiency of plaquing (EOP) tested for a panel of 10 phage. Darker colors indicate a higher level of protection. Systems leading to smaller plaque sizes are noted with an ‘S’ and systems that protect via an abortive infection (Abi) mechanism are indicated with an asterisk. For each new defense system identified, the operon structure and predicted domain composition of each component is shown. Shaded regions correspond to domain predictions using HHpred, summarized by association to PFAM clan, with short descriptions above. TM, transmembrane domain; HTH, helix-turn-helix. (**b**) Bacterial growth in the presence of phage at MOIs of 0, 0.05, or 5. Robust growth at MOI 0.05, but not MOI 5, indicates an Abi mechanism. See Fig. S3 for extended MOI data. (**c**) Plaquing of T4 on *E. coli* isolates ECOR22 and ECOR65 or the isogenic defense system deletions. Dilutions were done on two different plates and images combined for presentation. (**d**) Plaquing of T4 on strains harboring the indicated defense system or isogenic site-directed mutants of predicted domains. Asterisks indicate approximate location of mutations made. (**e**) Instances of homologs of defense systems by bacterial class, sorted by number of instances, descending from left to right. Known systems are listed in bold for comparison to newly identified systems.

We then sought to classify whether each system functioned via Abi or provided direct immunity such that infected cells could survive an infection. Because Abi systems require sacrifice of infected cells, these systems typically only provide defense at a low multiplicity of infection (MOI), in which bacteria outnumber phages, whereas direct immunity provides defense to the infected cell and thus allows comparable, though not identical, rates of growth at MOIs above and below 1.0. We thus tested the growth of strains harboring each defense system infected at an MOI of 0.05 and 5. Of the 21 systems tested, 9 provided direct inhibition of phage infection, producing comparable protection at both MOIs; the other 12 likely use an Abi mechanism of protection with stronger protection at the lower MOI (Fig. 3b, S3a). These results validate the ability of our screening strategy to detect Abi defenses and underscores the notion that Abi systems contribute considerably to *E. coli* immunity.

One advantage to our screen is that we have the strains from which these defense systems originated, in contrast to computational screens that have identified many defense systems from species that have sequenced genomes but are not readily available. We were thus able to delete candidate systems in their originating strains and test whether, in their native context, they protect against phage. Specifically, we tested the native role of PD-T4-2 and PD-T4-9, which originate from ECOR65 and ECOR22, respectively, strains to which T4 can adsorb but not infect. Deleting each system dramatically increased the plaquing efficiency of T4, demonstrating that these systems provide defense in both their original, native context and when introduced into *E. coli* K12 (Fig. 3c). We also asked whether our defense systems work in *E. coli* strains other than MG1655 by testing four candidate defense systems in ECOR13 and *E. coli* C, which are natively susceptible to the three phages. All four systems provided protection indicating that the function of the systems identified is not strictly dependent on strain background (Fig. S3b).

In total, 26 of 32 proteins in the 21 systems identified here were annotated in GenBank as either “hypothetical protein” or as containing domains of unknown function and had no primary sequence homology to any characterized anti-phage defense system. To more sensitively characterize each protein, we used HHpred to detect even remote similarity to PFAM domains^19^. This did not reveal any homology to a known system, and in the majority of cases, most of each protein remained uncharacterized. We were, however, able to detect potential similarity to some motifs or domains characteristic of defense systems *e*.*g*. nucleic acid binding or cleavage domains (Fig. 3a, Table S3). In the majority of cases this was limited to small regions hinting at enzymatic function, but not a mechanism of activation or specific targets.

Remote homology detection revealed several intriguing features uncharacteristic of known defense systems. This included (i) similarity to a ribosome-dependent ribonuclease (RelE) in conjunction with a phage-sheath-like domain, (ii) a three gene operon encoding an exotoxin A-like domain known to participate in bacterial virulence though not phage defense, an unknown protein and a SecB-like chaperone, (iii) a putative membrane-anchored protein with a central coiled-coil domain (DUF4041) and a C-terminal DNA binding/cleavage domain, (iv) a beta-propeller fused to a DNA binding/cleavage domain, (v) a P4 phage β-like protein, (vi) a lipoprotein, (vii) a zinc-finger like domain fused with a C-terminal domain of unknown function that belongs to an extended family of ribonucleases, (viii) a CoiA domain, (ix) and DUF6575.

Several defense systems showed similarity to domains that are less frequently associated with defense systems compared to domains such as nucleases and helicases. These less frequent domains included a peptidase, a eukaryotic-like Ser/Thr kinase^20^, a NAD^+^-binding Sir2 homolog^21^, and a GIY-YIG nuclease^22^. Four of the systems identified contained a component with similarity to a toxin of toxin-antitoxin (TA) systems, but either none of these systems were found in existing TA databases or they encoded additional uncharacterized components, aside from simply toxin and antitoxin. Collectively, our results reveal a striking diversity of proteins involved in bacterial defense and highlight the vast, unexplored landscape of antiviral immunity in bacteria.

In total, 11 defense systems had a component suggestive of DNA binding or cleavage activity (often including distant similarity to the restriction endonuclease-like fold PD(D/E)XK), though this similarity was, as noted above, typically restricted to a small region or motif. These putative DNases are unlikely to be part of restriction-modification systems as only one included a predicted methylase. Notably, 7 of 11 of these provided direct immunity (not Abi), suggesting non-self, nucleic acid-targeting activity in potentially novel ways. Remote similarity to HEPN motifs or domains were found in six of the 21 systems. These domains are also present in the ribonucleases associated with CRISPR-Cas and toxin-antitoxin systems, supporting the notion that they are common, versatile components of defense systems^23^. In four cases, we mutagenized key, conserved residues in the predicted domains and found they were essential to phage protection, suggesting the remote domain predictions feature in the function of these systems (Fig. 3d).

To assess the conservation of the 21 systems identified, we investigated the phylogenetic distribution of each (Fig. 3e). Homologs of each system (*i*.*e*. those that encode all components) were found in other gamma-proteobacteria, with 16 and 18 having homologs in alpha-and beta-proteobacteria, respectively. More than half were also abundant in Firmicutes, Actinobacteria, Bacteroidetes, and Spirochaetia, suggesting that many of the systems represent new, widespread classes of anti-phage defense systems. Seven of the systems we identified were restricted to proteobacteria and three were exclusively found in γ-proteobacteria. Thus, the phage immune landscape of *E. coli* is composed of both widespread and clade-specific systems.

### Mobile elements dominate the defense system landscape in *E. coli*

Prior searches for proteins enriched in defense islands identified and validated 38 novel defense systems^12,13^. None of the 21 systems identified here are homologs of those systems, and only one component (of PD-T7-2) resembles (32% identity) a protein of a previously validated multi-component system. PD-T4-8 has a DUF4263 domain in common with the Shedu defense system, but is not homologous to the validated *B. cereus* system. The computational approach in ref^13^ also identified 7,472 protein families enriched in defense islands that have yet to be validated. Only 14 of 32 proteins identified here have homology to those, and often with < 35% identity over limited regions of the proteins (Table S4). These observations suggested that our experimental-based selection may uncover different types of defense systems than can be found computationally by searching for enrichment in defense-islands.

To further probe this idea, we analyzed the native genomic context of the 21 systems identified here. We found that 12 of the 21 were located in intact prophages (Fig. S4). Seven of these systems were found within P2-like prophages, with four located in the same position of the P2 genome, directly between the genes encoding the P2 replication endonuclease and portal protein (Fig. 4a). This location has been previously found to harbor the defense systems *tin* and *old*, the former of which was also identified in our screen against λ_vir_^24^ (Table S1). More recently, this location was found to encode a wide array of previously uncharacterized defense systems^25^. We also observed a second defense-enriched locus in P2-like phages, which contained three of the systems discovered here, and the previously identified defense gene *fun* in the P2 reference genome^24^ (Fig. 4a). This hotspot was not noted in the recent analysis of P2 defenses. Notably, 47 of our 71 *E. coli* strains together encode 63 P2 portal proteins, with 110 unique proteins found in the adjacent hotspot. Only 13 of these 63 hotspots contain previously known defense systems, and most genes are annotated as hypothetical. These findings suggest than not only do P2 prophages encode a rich diversity of anti-phage proteins^25^, but that P2 defense hotspots make up a significant fraction of the immune landscape in *E. coli* (Fig. 4b).

**Figure 4.**
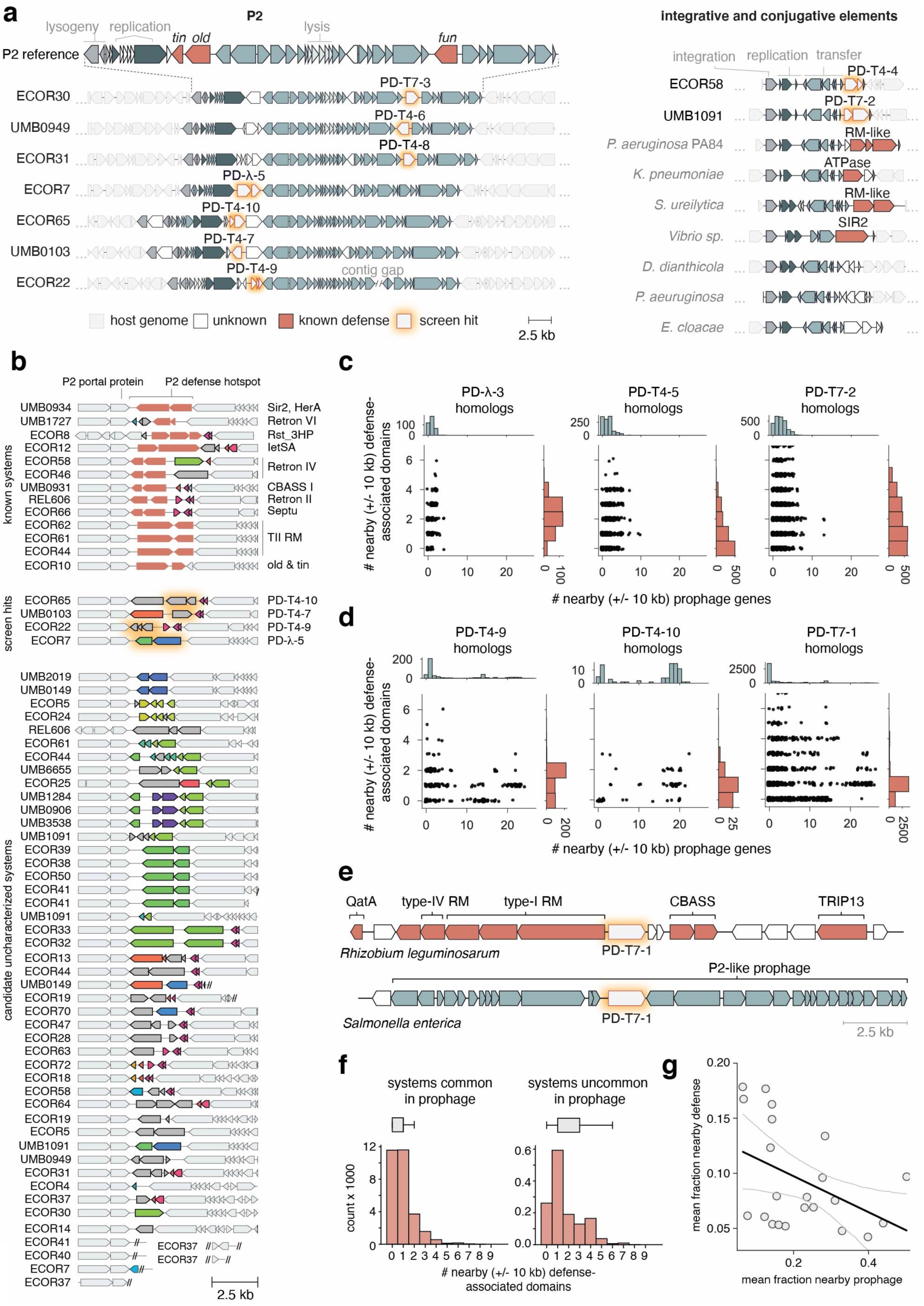
Prophages and mobile genetic elements are major sources of defense systems. (**a**) Hotspots of novel defense systems. (*Left*) The native genome context of 7 defense systems identified here, showing the boundaries of P2-like prophages in the genome from which they originated. Genes are color-coded as indicated below. (Right) Two defense systems were identified in the accessory region of an integrative and conjugative element (ICE) within the indicated *E. coli* genome. Homologous ICEs from other bacterial genomes contain known and putative defense systems in the same location. (**b**) All identified instances of P2 defense hotspot #1 in our 73-strain *E. coli* collection. White genes are flanking, conserved P2 genes. Each color of gene within the hotspot represents a protein cluster (30% identity). All grey genes belong to a lone cluster. Double slashes denote the end of a contig. (**c**) Number of defense and prophage-associated genes +/- 10 kb from system homologs. The scatterplots indicate, for each homologous system, the numbers of prophage and defense-associated genes within +/- 10 kb. Examples in (**c**) represent systems that were found outside of prophages in our genome collection. For all 21 systems, see Fig. S5. (**d**) Same as (**c**) but for systems we found in prophages. (**e**) Examples of the +/- 10 kb context for PD-T7-1 homologs in a defense island or prophage. (**f**) Distribution of number of nearby defense-domain containing genes in homologs of systems commonly found in prophages (>10% homolog-containing regions with 8+ prophage genes in proximity) or not; n = 12 and 9 systems, respectively. (**g**) Linear regression fit of total nearby prophage gene and defense-domain-containing genes for each system, Pearson r = -0.442, P = 0.045, error indicates 95% CI.

Five defense systems were found associated with other types of prophages or their remnants, including P4 satellite prophages or related integrases, Mu-like phages, and lambdoid phages (Fig. S4). We also observed a defense-enriched locus within an integrative and conjugative element (ICE), containing two systems we identified, PD-T4-4 and PD-T7-2 (Fig. 4a). This type of ICE is not widely distributed in our *E. coli* strain collection, but instances of this ICE in other genomes encode known defense systems at this locus, as well as hypothetical proteins that may also function in phage defense (Fig. 4a). PD-T7-4 and its homologs often overlap an integrase gene, while PD-T4-5 was identified on a plasmid. The remaining 4 systems did not appear to reside within active MGEs, but each had a nearby integrase gene suggesting they may be part of decaying MGEs (Fig. S4).

Our study supports prior findings that in addition to defense islands, prophages and other mobile genetic elements (MGEs) are a rich reservoir of defense systems^26^. However, these categories are not mutually exclusive, as some defense islands may be carried on or derived from MGEs^27^. To more systematically document the different genomic contexts for the systems identified here, we collected, for each of the 21 systems, all homologs in a set of 844,603 publicly available bacterial genomes. We classified the genes within 10 kb upstream and downstream of each homolog as either defense-related, prophage-related, or neither (see Methods). We then tabulated the number of defense-and prophage-related genes flanking each homolog (Fig. 4c, S5).

We detected two distinct patterns. For systems that we found outside of prophages in our strains, the homologs were also typically not prophage-associated (Fig. 4c, S5) and were often near several other genes encoding defense-associated domains. Thus, these systems do appear in defense islands, even though they were not previously detected as enriched in them. For the systems we identified within *E. coli* prophages, some of their homologs were also found in prophages, as evidenced by dozens of flanking prophage-related genes (Fig. 4d, S5). These prophage-associated homologs were typically near 1-2 defense-associated genes, but rarely more than 2, suggesting that some systems reside in small defense islands, or clusters, within a prophage, as with the P2 hotspots (Fig. 4d)^25^.

Notably, there are homologs of each system we identified that can be found in defense islands (Fig. 4e), some more rarely than others, indicating that they do not require a prophage context to function. In aggregate, we observed an inverse correlation between the number of defense-associated genes for homologs in prophages versus not (Fig. 4f-g). This highlights the constraint on how many defense systems can be carried by prophages, or within a given hotspot, due to size limitations in DNA that can be packaged. We suspect that this is one reason why many of these genes are enriched in prophages as compared to defense islands.

Finally, we compared defense island and prophage enrichment (see Methods) between systems discovered here and those previously predicted computationally and experimentally validated^12,13^. We found that our experimentally selected systems on average were less frequently associated with known defense genes, but more frequently associated with prophage genes (Fig. S6). These analyses suggest that defense-island enrichment methods may be less sensitive in identifying defense systems frequently found in prophage.

### Novel toxin-antitoxin-like defense systems

Toxin-antitoxin (TA) systems are typically comprised of a protein toxin that can arrest cell growth, but is normally neutralized by a cognate, co-expressed antitoxin. TA systems are extremely prevalent in bacterial genomes and MGEs, but their functions remain poorly understood^28^. A handful of TA systems have been found to provide anti-phage defense through an Abi mechanism^29,30,30,31^. Our selection yielded four different systems that were recognizable as TA-like in nature. These encoded gene products with sequence similarity to toxic proteins, mostly featured multiple components, and provided Abi defense (PD-T4-5, PD-T4-7, PD-T4-9, PD-λ-2). A fifth (PD-T4-10) facilitated Abi defense and had two overlapping ORFs, reminiscent of many TA systems. As noted above, none of these systems were previously annotated as TA systems so we sought to validate the three featuring multiple components: PD-T4-10, PD-λ-2, and PD-T4-9 (Fig. 5a).

**Figure 5.**
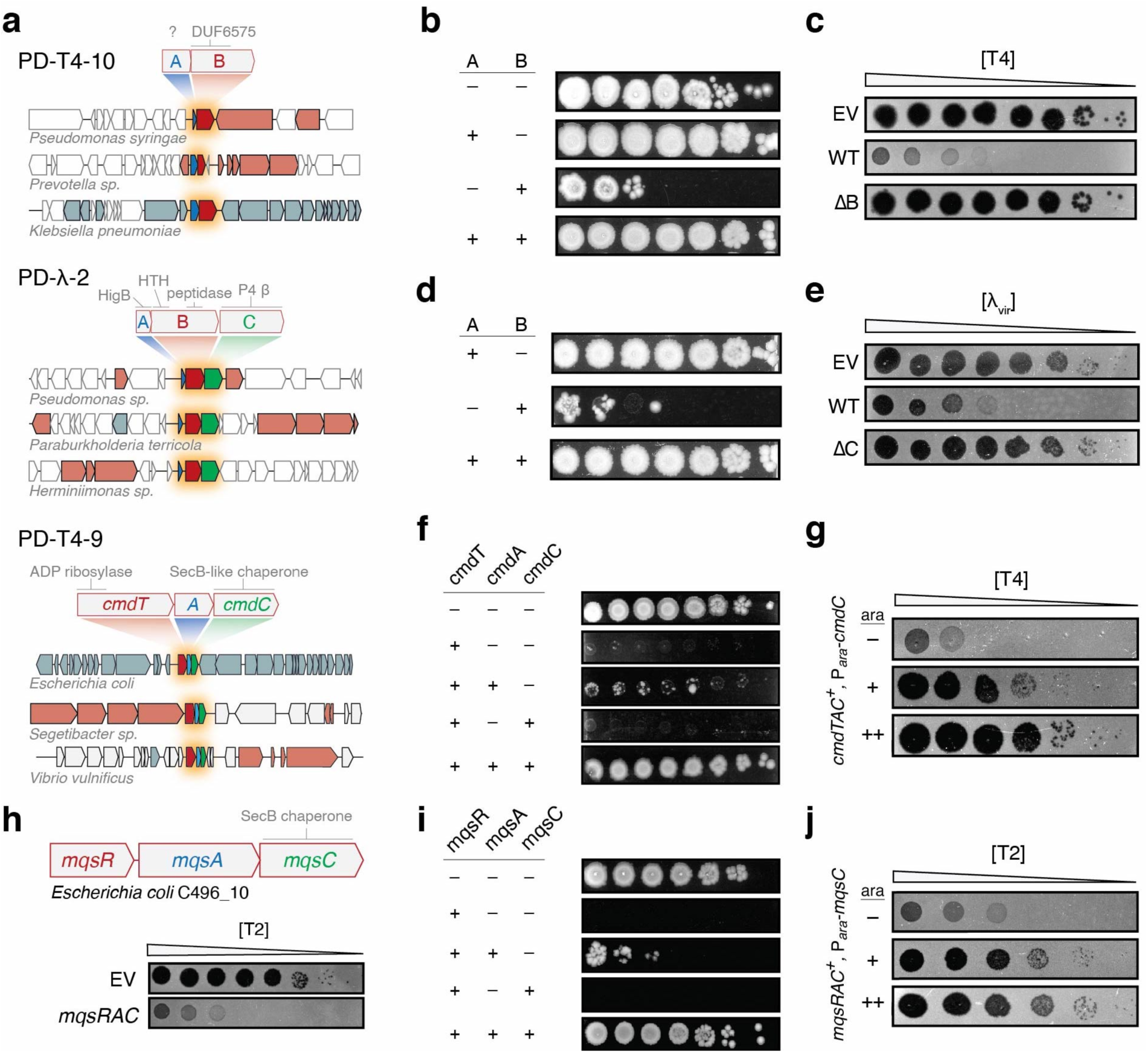
Novel toxin-antitoxin derived defense systems. (**a**) Schematics of PD-T4-9, PD-T4-10, and PD-λ-2 defense system operons and their domain predictions. Representative homologs of the systems are shown in their genomic contexts and indicate conservation and order of the system components. Blue, putative antitoxin; red, putative toxin; green, accessory factor. (**b, d, f, i**) Each component or pair of components indicated was expressed (+) or not (-) from an inducible promoter and assayed for viable colony-forming units in 10-fold serial dilutions. (**c, e**) Plaquing assays for the phage indicated on cells harboring an empty vector or a vector containing a given defense system with all components (WT) or lacking the component indicated. (**g, j**) Plaquing of phages on TAC-containing strains expressing a second copy of the chaperone component to varying levels during infection. (**h**) Schematic of *mqsRAC* system.

For PD-T4-10, neither of the two proteins had a predicted function. We expressed each component from an inducible promoter and found that the second component, PD-T4-10B, was toxic. This toxicity was fully neutralized when PD-T4-10A was co-expressed (Fig. 5b). PD-T4-10A could not be deleted, consistent with it being an antitoxin, whereas deletion of the toxin PD-T4-10B abolished resistance to T4 infection (Fig. 5c). Thus, this system comprises a novel, *bona fide* TA system that provides strong protection against T-even phages.

PD-λ-2 features three components. The first is similar to HigB toxins that inhibit translation, the second has a HigA antitoxin-like domain (Xre-family helix-turn-helix) fused to a C-terminal peptidase domain, and the third component is related to a P4 phage antitoxin. An Xre-peptidase fusion, co-expressed with an upstream toxin-like gene, has been observed in sequence-based TA searches but the function of these loci was unknown^32^. Unexpectedly, we found that overexpressing the second component, PD-λ-2B, alone was toxic, with toxicity rescued by co-expression with PD-λ-2A (Fig. 5d). We also confirmed that PD-λ-2C, though not required to neutralize the toxin, is required for defense against Ψ_vir_ (Fig. 5e).

PD-T4-9 also contains a third component, a SecB-like chaperone, suggesting that it is related to an enigmatic class of TA systems called toxin-antitoxin-chaperone (TAC) systems so we renamed this system CmdTAC for <u>c</u>haperone-<u>m</u>ediated <u>d</u>efense. The antitoxins of TAC systems are typically homologous to canonical antitoxins, but feature an unstructured, hydrophobic extension of their C-termini called the chaperone addiction domain (ChAD)^33^. In the absence of the cognate SecB-like chaperone, the ChAD renders the antitoxin prone to aggregation and proteolytic degradation, thereby freeing the toxin^33^. Inducing the expression of CmdT, which has an ADP-ribosyltransferase-like domain, was toxic. Co-expression with the presumed antitoxin, CmdA, only marginally improved viability. However, co-expression with both the putative antitoxin and chaperone components completely restored viability (Fig. 5f). Together, these results suggest that CmdTAC is a novel TAC system that protects against phage.

To further characterize the role of the chaperone, CmdC, we overproduced it during T4 infection of CmdTAC^+^ cells. Interestingly, an oversupply of CmdC abolished phage protection by CmdTAC, suggesting that destruction of CmdC, or sequestration of CmdC from the complex may allow this TA system to activate in response to phage infection (Fig. 5g). The chaperone normally promotes neutralization of CmdT by CmdA, but, following infection, could be depleted or sequestered by a phage product, leading to liberation of the toxin and abortive infection. Providing additional CmdC prevents the loss or complete sequestration of chaperone, and thereby prevents release of CmdT from CmdA. Future study is ongoing to further dissect the mechanistic nature of chaperone-phage interactions.

We hypothesized that TAC systems may be a broad class of phage defense systems. To test if other TAC systems can protect *E. coli* MG1655 against phage infection, we cloned and tested an MqsRAC system from *E. coli* C496_10. Although completely unrelated in toxin and antitoxin sequence to CmdTAC, MqsRAC is a canonical TAC system, has been characterized in Mycobacteria, and includes a SecB-like chaperone homologous to CmdC^33^. This system conferred robust protection against T2 (Fig. 5h), but not T4 as with *cmdTAC*. Like CmdTAC, toxicity of MqsR could only be rescued by expressing MqsA and MqsC (Fig. 5i), and inducing additional MqsC in MqsRAC^+^ cells inhibited phage protection (Fig. 5j). Thus, our work indicates that TAC systems may be a widespread and diverse new class of phage defense system.

## Discussion

Our work indicates that a large reservoir of diverse, previously unknown phage defense genes is distributed across the *E. coli* pangenome. Like many bacteria, there is tremendous variability in the ‘accessory’ genomes of different strains of *E. coli*. Many of these accessory genes are likely associated with phage defense, as recently suggested for marine *Vibrio*^34^. Although efforts to find new defense systems based on proximity to known systems have proven fruitful, our work reveals that many phage protective systems remain unidentified.

Our functional screening, which is agnostic to the genomic context of defense systems, indicates that many systems may not be commonly or detectably enriched among known defense islands. Indeed, none of the 21 systems we identified were previously reported to provide phage defense in prior discovery efforts based on defense island enrichment. Moreover, only 3 of the systems we identified were within 10 kb of other known defense systems in our *E. coli* genomes. However, notably, 15 of the 21 systems we identified were present in apparently active or recently active MGEs (prophages, ICEs, or plasmids) (Fig. S4), with the other 6 located in regions suggestive of MGEs in the late stages of decay. Homologs of the systems discovered here are sometimes present in defense islands (see Figs. 4d, 5a), but these associations are often relatively rare.

It is well established that MGEs contribute to antiviral defense in bacteria^26^. By providing a glimpse into the relative contributions of MGEs and defense islands to immune system context, our results support the notion that active prophages and other MGEs are likely the primary reservoirs of anti-phage defense systems in *E. coli*. This idea is consistent with studies that have identified diverse anti-phage systems in P2 and P4 prophages^24,25^ and another recent study revealing how the transfer of defense-containing ICEs drove the emergence of phage resistance in clinical isolates of *V. cholerae*^34^. The defense systems found in functional MGEs likely help these elements to protect themselves and host resources by preventing infection by other phages^27^. However, given genome size and packaging constraints, there is a limit to the number of defense systems carried by a given prophage. Such constraints may help explain why increased prophage-association was correlated with lower defense gene association (Fig. 4g). Future investigation should work toward further uncovering the vast reservoir of anti-phage elements carried on MGEs.

Of the 32 proteins in 21 systems found here, 13 feature domains never before reported to function in phage defense. These include a protein with distant similarity to RelE and a region of a phage tail sheath, an Exotoxin A-like domain previously only shown to function in bacterial virulence, a SecB-like chaperone, DUF6575, DUF4041, a CoiA competence-related domain, a β-propeller fold, an ImmA/IrrE peptidase, a HigB toxin, and three proteins with no ascertainable similarity to deposited domains. In addition, 10 proteins contained large regions with no predicted domains (Fig. 3a).

Although some regions of the proteins identified have distant similarity to known nuclease motifs, these domains are found here in new or unusual contexts and associations, which raises fascinating questions for future investigation. For example, one features a 7-bladed beta propeller with a separate N-terminal Mrr-family nuclease domain. The β-propeller fold consists of separate modules that adopt a disc-like, circularly arranged structure with a central channel that can accommodate many substrates including protein and DNA^35^. Determining how this domain aids in activation or target specificity of this system opens many avenues of future discovery. We also uncovered 5 single protein systems that exclusively contain a putative DNase domain. Although these domains are found in defense systems like RM and CRISPR, how the orphan DNase-like proteins sense and respond to phage infection, especially to provide direct defense, is unclear. As all but one is encoded without a DNA methylase, they likely do not distinguish phage and host DNA in the same way as RM systems. Additionally, these DNase-containing systems protected against T4 and T7, which are intrinsically resistant to most RM systems^36,37^. Type-IV RM systems are single component defenses known to target the modified cytosine of T-even phages^38^, but none of the 5 proteins discussed here resemble these. How those that are specific to T7 might target T7 DNA, which is not modified, is unclear. The existence of unconventional, phage-targeting, nucleic-acid degradation systems underscores a knowledge gap in the molecular mechanisms of viral resistance and self/non-self recognition.

We also identified and validated three TA systems as phage defense elements including *cmdTAC*, prompting the discovery that unrelated TAC systems such as *mqsRAC* also function in antiviral immunity. MqsRA is a well studied TA system, but with no documented role in phage defense; our results suggest that bacteria have co-opted this TA system by addition of a chaperone-dependent antitoxin in order to activate the toxin in response to infection. Our findings support the notion that TA systems play a central role in phage defense^29^.

Finally, we note that four systems identified here likely belong to classes of previously described systems, but have diverged quite significantly such that they share little significant sequence homology, reinforcing the extreme divergence and adaptations that typify many immune systems. These divergent systems include PD-T7-2, in which the second protein is similar to HerA of Sir2/HerA^13^, PD-T4-8, whose central domain bears distant resemblance to Shedu^12^, PD-λ-5, which appears to be a highly-compacted prophage version of an RM + Abi system, and PD-T4-5, a plasmid borne gene that is a distant relative of AbiF/D.

Our screening methodology enables the experimental discovery of anti-phage defense genes and has several powerful features. First, we can return to the strain of origin for a given system and demonstrate that it provides defense in its native context. Second, there is a built-in pairing of defense systems to the phage they defend against, whereas with computational studies, the phage(s) a given system defends against must be subsequently identified. Third, in the pipeline described here, the source DNA comes from other *E. coli* strains, which likely minimizes false negatives that can arise from producing candidates in a heterologous host. Finally, our experimental approach is not limited to genes that are detectably enriched in defense islands. As noted, only three of the systems we identified were natively associated with obvious defense islands and some also do not appear to have many close homologs in defense islands. Some defense systems may not associate with defense islands while some may have arisen too recently or not be widespread enough to detect an association. Indeed, some of the systems identified here show a relatively limited phylogenetic distribution. However, the phage defense capabilities of bacteria likely include both broadly conserved and clade-specific systems adapted to the unique biology of a given organism and its phages.

The methodology developed here can be powerfully extended in several ways. First, genomic DNA from other sources, including metagenomic DNA, could be used as input material. From just 71 strains we identified 21 new defense systems suggesting fertile ground remains for discovery both in and beyond *E. coli*. Second, the panel of phages tested here was limited to three and could easily be expanded, particularly given the enormous diversity of phages. Finally, with only small modifications, any transformable bacterium could be used as the host strain. Further identification and characterization of bacterial immune systems promises to shed new light on the ancient arms race between bacteria and their viral predators, and may also have practical applications, providing the foundation for precise molecular tools and helping to inform future efforts to develop phage as therapeutics.

## Materials and methods

### Bacteria and phage growth and culture conditions

Cultures were routinely grown at 37 °C in LB unless otherwise stated. Phage stocks were propagated on MG1655, filtered through a 0.2 μm filter, and stored at 4 °C. Select ECOR strains were obtained from the Thomas S. Whittam STEC Center at Michigan State University and UMB isolates were obtained from Alan J. Wolfe at Loyola University Chicago (Table S5). Other strains, plasmids, and primers + synthetic gene fragments are listed in Tables S6, S7, and S8, respectively.

### Library construction

Genomic DNA was harvested from pooled, overnight cultures of each *E. coli* isolate using the PureLink Genomic DNA Mini Kit (Invitrogen). From this sample, a fosmid library was constructed by Rx Biosciences Inc. (Gaithersburg, MD) using the CopyControl Fosmid Library Production Kit (Lucigen) according to the manufacturer’s protocol. Plasmid sub-libraries were constructed first by extracting fosmid DNA from select positive clones using the ZR Plasmid Miniprep Kit (Zymo Research). Equimolar, pooled fosmids were sheared to an average of 8 kb using g-TUBEs (Covaris). Sheared DNA was end-repaired and 5’-phosphorylated using the End-It DNA End-Repair Kit (Lucigen), then purified using the DNA Clean and Concentrator kit (Zymo Research). The plasmid vector was prepared by PCR and blunt-ended fragments were ligated to the plasmid using T4 DNA Ligase (NEB) for two hours at room temperature. The ligation reaction was electroporated into MegaX DH10B™ T1R Electrocomp™ Cells (Thermo Fisher) and selected on LB with 50 μg/ml kanamycin.

### Defense system selection

We used a variant of the previously described *tab* (T4 abortive) selection procedure to select for fosmids that provide resistance to phage infection^17^. A heavy inoculum (>30 μl) of a high-titer library freezer stock or empty vector freezer stock was inoculated into 5 ml LB containing 20 μg/ml chloramphenicol (Cm) and grown to stationary phase at 37 °C (approximately 4 hours, OD_600_ = 2-3). Cultures were adjusted to OD_600_ = 1.0 and 0.1 ml (∼8 × 10^7^ cells) was pipetted onto one side of 3-6 empty 15 cm Petri dishes. A 10-fold dilution series of phage stock was prepared, and 0.1 ml of each dilution was pipetted onto the empty plates containing the bacterial cultures (onto a separate area of the plate, preventing mixing). 20 ml of molten LB 0.5% agar were added to the plate and briefly mixed to disperse bacteria and phage. Plates were incubated at 37 °C overnight, with the exception of one T4 screen which was conducted at room temperature. Bacterial colonies were picked from the plate containing the phage dilution that produced the largest difference in number of colonies between the control and library samples, then streaked onto fresh plates to isolate single colonies. To test phage resistance phenotypes, single colonies were cultured overnight, and 30 μl of culture were mixed with 5 ml of molten LB 0.5% agar, 30 μg/ml Cm in 8-well rectangular dishes. Serial dilutions of phage were spotted onto the solidified culture media and incubated overnight at 37 °C. Clones with defense phenotypes, as described below, were stocked and miniprepped, with the ends of each fosmid insert Sanger sequenced.

With some phage-resistant clones we observed no lysis even at very high phage concentrations (see Fig. 2b). All sequenced clones with this phenotype contained either LPS or capsule biosynthesis genes. In our experience, no intracellular defense system completely prevents visible lysis at extremely high phage concentrations, whereas changes in the phage receptor or cell surface can, so we suspected that these clones disrupted phage adsorption. Similarly, with regard to T7, genes for capsule biosynthesis survived selection, but did not display any difference from the control in plaquing efficiency. All clones showing a complete lack of lysis, or no change in EOP and encoding capsule genes, were discarded. 12 positive clones (all from the Ψ_vir_ selection) produced a high number of discrete escape plaques consistent with RM systems. Strains that displayed this escape phenotype and whose fosmids contained an identifiable RM system were discarded from further analysis.

Sub-libraries were screened as described above using a variation which allowed bulk harvesting of all positive colonies directly from the screening plate (similar to the previously published *gro* screen)^39^. In this variation, instead of molten LB 0.5% agar, bacteria and phage samples were spread on the surface of LB 1.2% agar using glass beads.

### Long-read sequencing and defense system identification

After cells were harvested in bulk from the screening plates, total plasmid DNA was extracted and linearized by digestion with the restriction enzyme NdeI, EagI, or FsoI. For Oxford Nanopore sequencing, linearized samples were characterized on a FemtoPulse (Agilent Technology) to confirm integrity and high-quality samples were indexed by native barcoding (ONT kits EXP-NBD104/114) with supporting reagents from New England Biolabs. Libraries were prepared using the LSK-109 chemistry and samples were run on either a MinION (R9) or PromethION (R9.4) flowcell. Basecalling was done using built-in ONT tools. Processed reads were aligned to public reference genomes of the source organisms, or the relevant portions of the genomes contained in the fosmid inserts, using the Minimap2 plugin within the Geneious Prime 2020.2.4 software suite. Fosmids that could not be mapped to their genomes due to contig gaps were also sequenced identically, and *de novo* assembly was conducted using Flye^40^, also in Geneious. Candidate defense systems were predicted to be any gene or operon residing under the coverage maxima. In the minority of cases in which the result was ambiguous, candidates were cloned after prioritization by features including domain prediction, location in defense hotspots, hypothetical proteins, and by general comparative genomic investigation. Multi-component systems (operons) were predicted by ORF proximity, promoter prediction, and gene co-occurrence in homologs.

### Strain construction

Defense system cloning was performed using Gibson Assembly of PCR products containing predicted defense systems and their predicted promoters into a destination vector lacking an upstream promoter. Assembled plasmids were transformed into MG1655 by the TSS method^41^. MG1655 with a deletion of the region containing *mrr* was used as the host strain for PD-λ-5. Site-directed mutagenesis was conducted by PCR using outward facing primers containing the desired mutation and with compatible overlapping regions. Amplification of the wild-type template plasmid was cycled 18 times and the reaction was chemically transformed into DH5α cells. In-frame deletions of defense systems were constructed by transforming a temperature-sensitive plasmid expressing λ-red recombinase into the target strain. Oligos with overlapping regions to the genome targeted for deletion were used to amplify a kan^R^-resistance marker. The amplicon was then electroporated into the target strain induced to express λ-red and recombinants were selected on kanamycin. The recombinase plasmid was then cured from the target strain by growth at 37 °C.

### Efficiency of plaquing assays

50 μl of overnight cultures grown at 37 °C were mixed with 3 ml of LB 0.5% agar and overlayed onto LB 1.2% agar plates containing appropriate antibiotics. 2 μl of phage from a ten-fold serial dilution of stocks were pipetted onto the surface of the overlay plate. Spots were allowed to dry and incubated at 37 °C until plaques were visible. Plaques were then enumerated and EOP was measured as total plaque forming units (PFU) on the experimental strain divided by PFU on the control WT strain. Often, individual plaques were not distinguishable, *i*.*e*. no viable phage were produced in an infection, resulting in a lysis zone but no discrete plaques. In such an event, samples were counted as having one plaque on the last dilution that showed lysis. For EOP assays with TAC systems, chaperone expression was titrated by overlaying cultures on media with increasing concentrations of arabinose before spotting phage dilutions.

### Bacteriophage adsorption assay

Method is adapted from ref^42^. Overnight bacterial cultures were diluted 1:100 and grown to OD_600_ = 0.5. Cultures were infected at an MOI of 0.1. Samples were then incubated at 37 °C (T4 and λ_vir_) or 25 °C (T7) for 15, 25, or 15 minutes for T4, λ_vir_, and T7, respectively. 500 μl samples were then added to a tube of ice-cold chloroform, vortexed, and unadsorbed phage were enumerated by the top agar overlay method using a susceptible indicator strain. Percent adsorption was determined relative to a simultaneous mock control experiment that contained growth medium but no host cells.

### Abortive infection assays

Overnight cultures were normalized to OD_600_ = 1.0 and then diluted 100-fold. 150 μl of diluted cultures were dispensed in a flat-bottomed 96-well plate. 10 μl of phage dilutions were added to each well such that the MOI varied from 50 to 0.005. Wells were then overlayed with 20 μl mineral oil and plates were covered with a breathable membrane. Plates were incubated at 37 °C in a Biotek Synergy H1 Microplate Reader. OD_600_ was measured every 15 minutes. Three technical replicates were conducted for each strain.

### Toxicity assays

Strains containing plasmids with inducible promoters were grown overnight at 37 °C in LB under repressing conditions (LB or LB + 0.2% glucose). Cultures were washed in LB and ten-fold serial dilutions were spotted on LB agar with and without inducer (0.2% arabinose, 200 μg/ml vanillate, or 100 μg/ml anhydrotetracycline). Plates were incubated overnight at 37 °C.

### Bioinformatic analyses

Sanger sequences of fosmid ends were mapped to their strains of origin using BLASTn^43^ followed by manual inspection. General remote domain prediction was done using the HHpred online web server (https://toolkit.tuebingen.mpg.de/) or locally (HHblits and HHsearch) against Pfam A domains (v. 35.0)^19,44^. To label domains in Figure 3, we used the top HHsearch hit for each independent region of the protein. If there were many good matches, the bounds of the predicted domain were taken from the top hit, while the label was chosen based on the Pfam clan to which the top hits belonged. The only exception was PD-λ-5, for which the top hit, “methyltransferase”, was chosen as we deemed it more descriptive than the Pfam clan designation. Investigation of the P2 defense hotspot was conducted by identifying homologs of P2 portal protein using BLASTp, extracting the surrounding genes, clustering to 30% protein identity and visualizing using Clinker and clustermap.js^45^. Pan-genome analyses were performed by annotating Genbank assemblies with Prokka^46^ followed by analysis with Roary ^47^. The phylogenetic tree was generated using FastTree^48^ on the core genome alignment produced by Roary, using a generalized time-reversible model. Other software that was instrumental for routine genome analyses were PATRIC webserver, DNAFeaturesViewer, and Mauve^49–51^.

To assess whether defense systems were potential toxin-antitoxin systems, we used WU-BLAST 2.0 to search against TADB v2.0 (https://bioinfo-mml.sjtu.edu.cn/cgi-bin/TADB2/nph-blast-TADB.pl)^52^.

### Identification of defense system homologs and genomic context analysis

For each defense system, we searched for homologs of each individual component using blastp against all bacterial proteins in the NCBI non-redundant (nr) protein database using the following parameters: -evalue 0.00001 -qcov_hsp_perc 80 for single-gene systems and -evalue 0.00001 for multi-gene systems. The NCBI nr database was downloaded for local use in March 2021. All instances of the homologs identified in the nr search were then located within all full bacterial genomes (n = 844,603) downloaded from Genbank and RefSeq (ftp://ftp.ncbi.nlm.nih.gov/genomes/all/) in April 2021. For multi-gene systems, the system was only considered present in a given genome if all components of the system were present in the same genomic region. The one exception was CmdTAC in which otherwise clearly homologous systems were widely variable in the antitoxin sequence. For this system, homologs were required to have a CmdT homolog, a second downstream ORF, and a SecB-like chaperone as the third component.

The local genomic context of a defense system homolog was defined as all coding sequences ± 10 kb of the system (or to the end of a contig if less than 10 kb). All coding sequences within this local context were searched for defense-related and prophage-related domains using HMMER3 hmmscan^53^ with E-value cutoffs of 10^−5^ and 10^−15^ for the defense-related and prophage-related domain searches, respectively. For the defense-related domain search, sequences were searched against defense-related pfam and COG domains identified and used in Makarova *et al*.^11^, Doron *et al*.^12^, and Gao *et al*.^13^. We considered each gene flanking a given homolog separately, even if multiple, adjacent genes were part of a single, multi-component defense system. For the prophage-related domain search, sequences were searched against all pVOG^54^ domains available as of May 2021 when the pVOG database was downloaded. For scatterplots and marginal histograms in Figures 4 and S5, any regions with < 10 coding sequences (*i*.*e*. located on short contigs) were excluded. In native context schematics, prophage genes were predicted by annotation, pVOG analysis and by BLASTp against the ACLAME database^55^.

To compare defense and prophage context between our systems and those that were identified computationally and subsequently validated in Doron *et al*.^12^ and Gao *et al*.^13^ (see Fig. S6), we identified homologs of each system and their flanking genes as described above. The collected flanking proteins were then clustered using the function cluster within MMseqs2^56^ with the following parameters: --cluster-mode 1 --min-seq-id 0.9. Each resulting cluster was called as defense-or prophage-related if at least 90% of the proteins within the cluster contained the same defense-or prophage-related domain(s), respectively. This clustering helps to control for overrepresentation of closely related sequences. Defense and prophage enrichment for a given system was then calculated as the number of defense or prophage domain containing clusters divided by the total number of clusters.

### Taxonomy analysis

The taxonomic distribution of each system was defined by the system’s presence across the downloaded bacterial genomes with the same parameters as described above. For a given genome with a defense system present, the NCBI taxid was extracted and translated to major bacterial classes using taxon kit^57^. For comparison, we also examined the taxonomic distribution of the following known systems: type I-IV RM systems, EcoKI, EcoRI, EcoPI, and McrBC, respectively; P2 old, AAD03309.1; Cas9, WP_032462936.1; Zorya I, system containing BV17222.1; Zorya II, system containing ACA79490.1; ToxN, WP_000675353.1; Kiwa, system containing AEZ43441.1; Druantia, system containing ERA40829.1.

### Comparison of novel defense system datasets

To assess whether the defense systems found here were present in the defense system dataset from Gao *et al*.^13^, we extracted the available representative protein sequences from the supplied tables in Gao *et al*. and clustered them with sequences identified here using MMseqs2^56^ at > 20% identity and > 50% coverage thresholds. If a protein sequence formed an independent cluster, we called it as absent from their dataset. In addition, we used DefenseFinder^58^ on our protein sequences to confirm that homology to known systems could not be detected.

## Acknowledgements

We thank I. Frumkin, C. Guegler, M. LeRoux, S. Srikant, and T. Zhang for comments on the manuscript.

## Additional information

Supplementary Information is available for this paper. Correspondence and requests for materials should be addressed to ML.

All data generated or analyzed during this study are included in the published article (and its supplementary information files). Code is available upon reasonable request.

---

**Figure S1.**
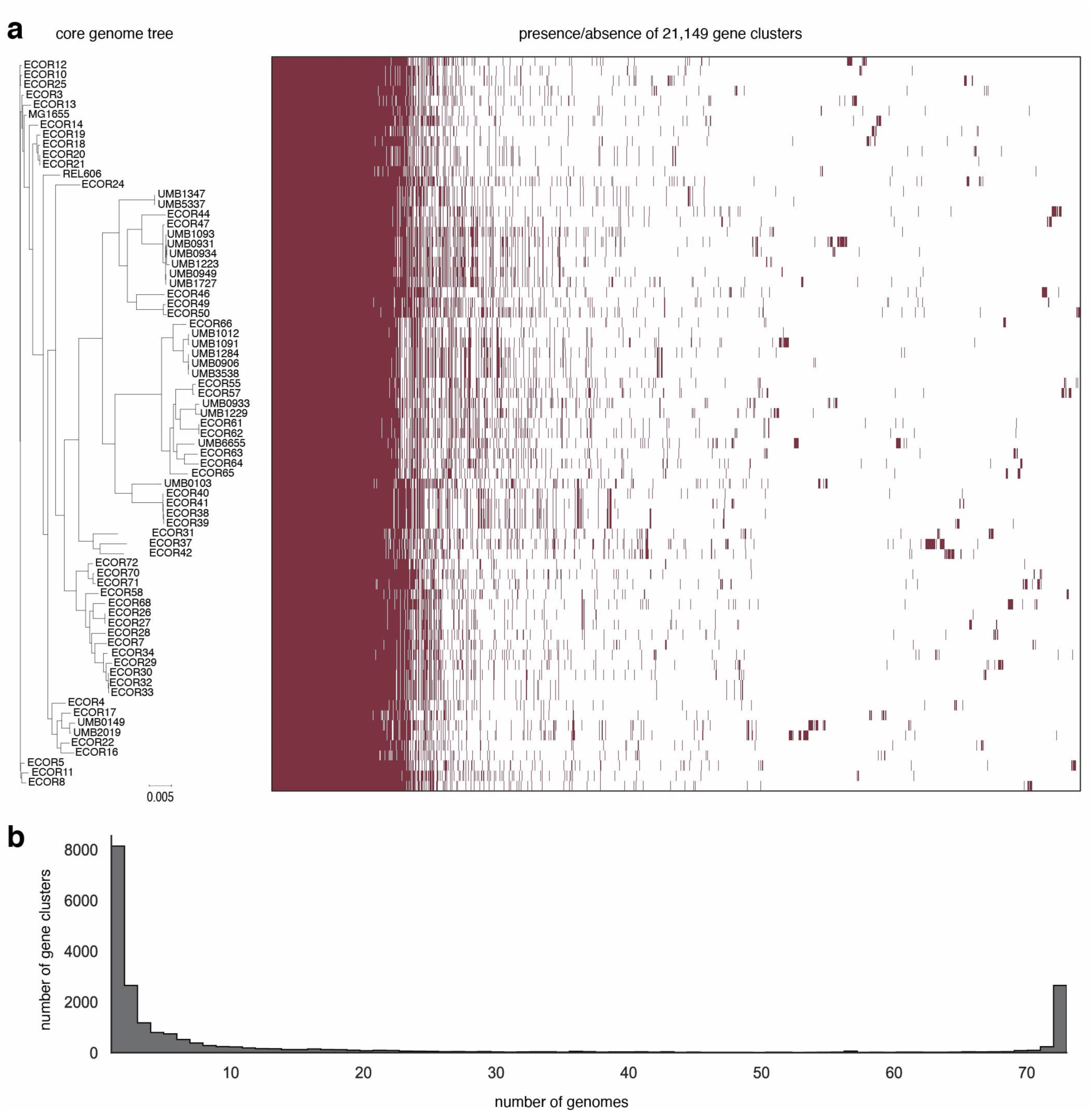
Diversity of *E. coli* isolate pangenome used in this study. (**a**) (*Left*) Phylogenetic tree of *E. coli* strain collection used to construct the genomic library screened. *E. coli* K-12 (MG1655) and B (REL606) are also included. (*Right*) Bars indicate presence/absence (red/white) of individual gene clusters (95% identity threshold). (**b**) Plot of the number of gene clusters versus the number of strains they are found in, *e*.*g*. ∼8,000 clusters are each found in only one genome. These sparsely conserved clusters represent the accessory genome, whereas ∼3,000 clusters are found in all 73 genomes and represent the *E. coli* core genome.

**Figure S2.**
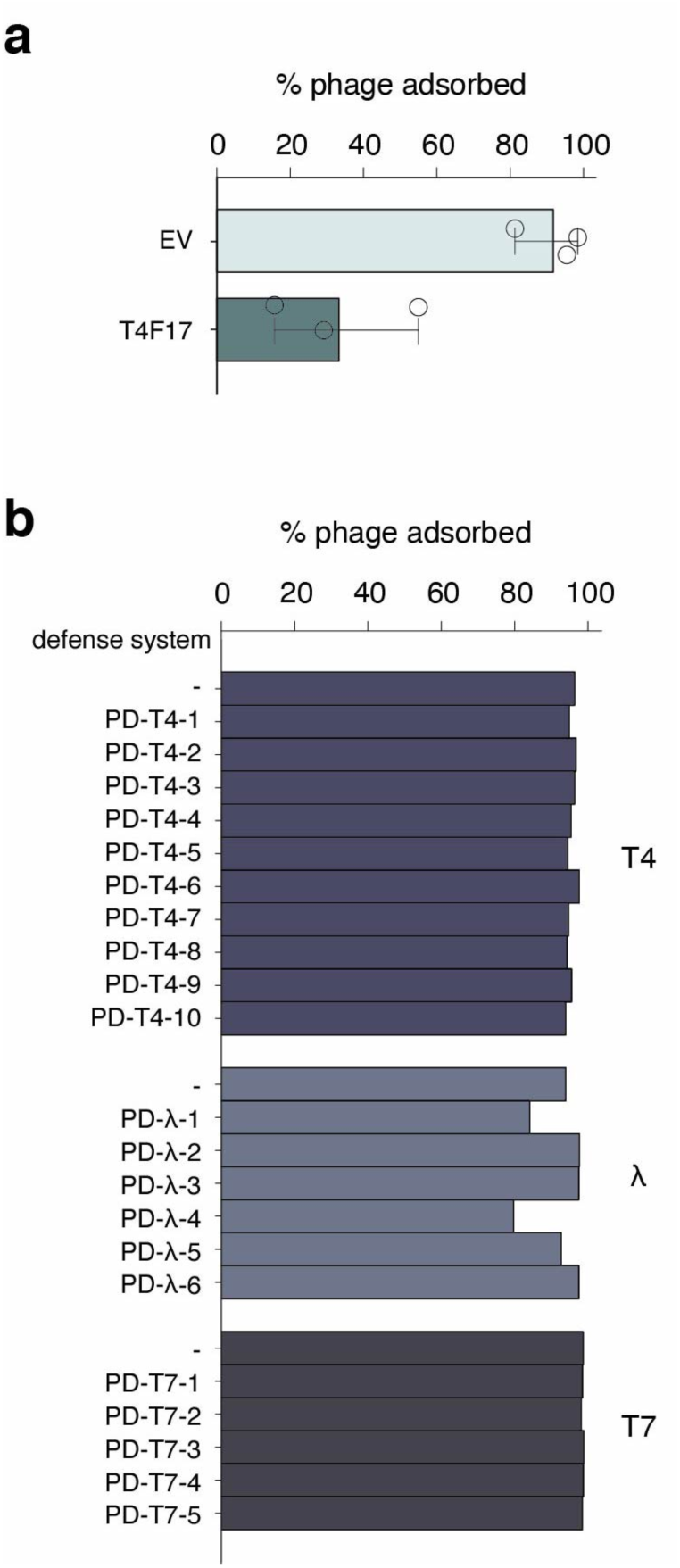
Adsorption of bacteriophage on various strains. (**a**) Adsorption of T4 on control (EV) and LPS-containing fosmid strains (T4F17). Error bars represent standard deviation of three biological replicates. (**b**) Adsorption of T4, λ_vir_, or T7 on strains expressing defense systems against their respective phage.

**Figure S3.**
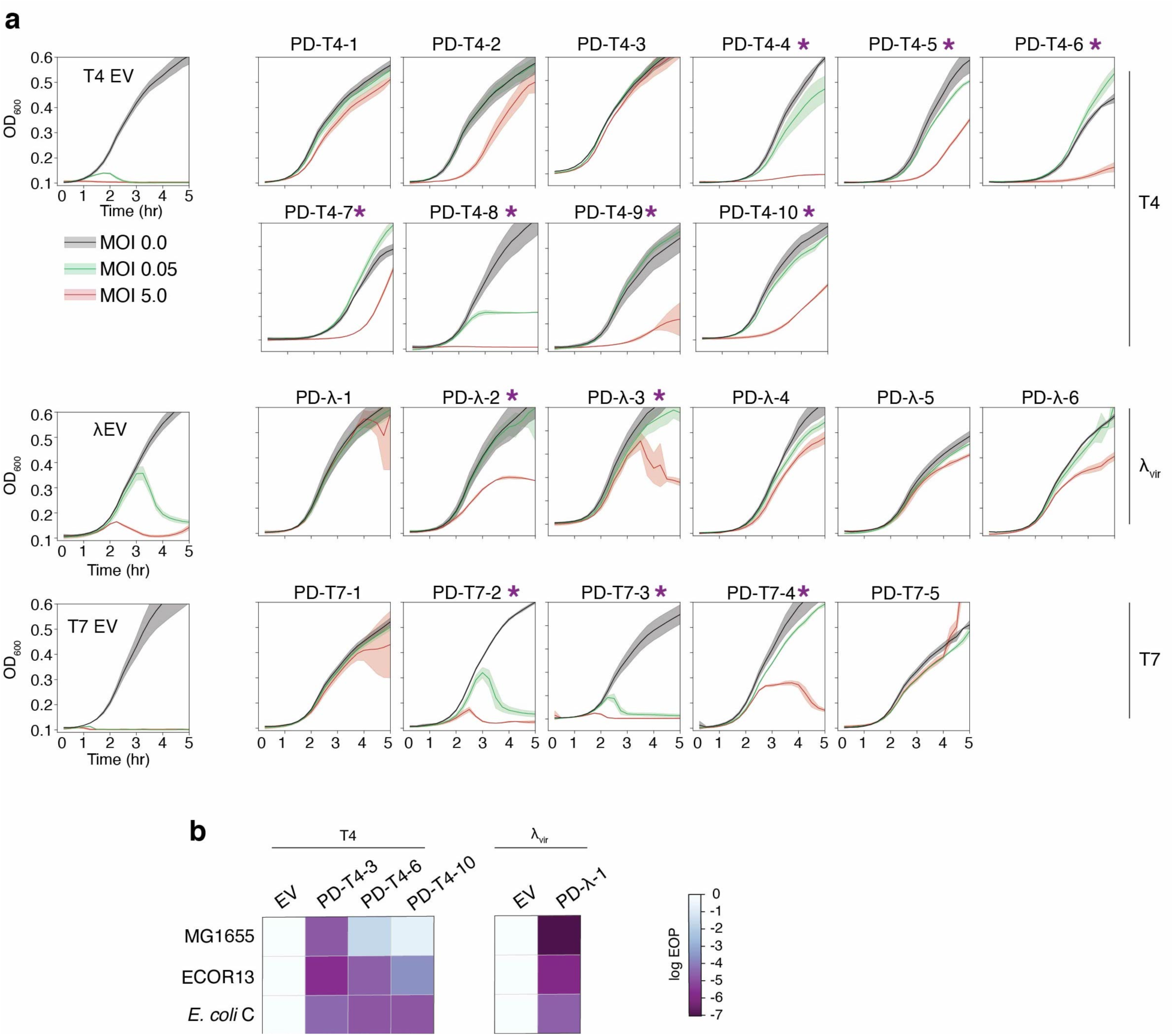
Abi mechanisms of defense and novel defense systems in other strains. (**a**) Growth of control (EV) or defense system-expressing strains with phage MOIs of 0, 0.05 or 5. Phages used in each experiment are shown on the right. Asterisks indicate systems not showing direct immunity and likely representing abortive infection mechanisms. Data represent three technical replicates with shaded regions indicating standard deviation. (**b**) EOP measurements for T4 (top) and λ_vir_ (bottom) on *E. coli* strains MG1655, ECOR13, or C expressing various defense systems indicated.

**Figure S4.**
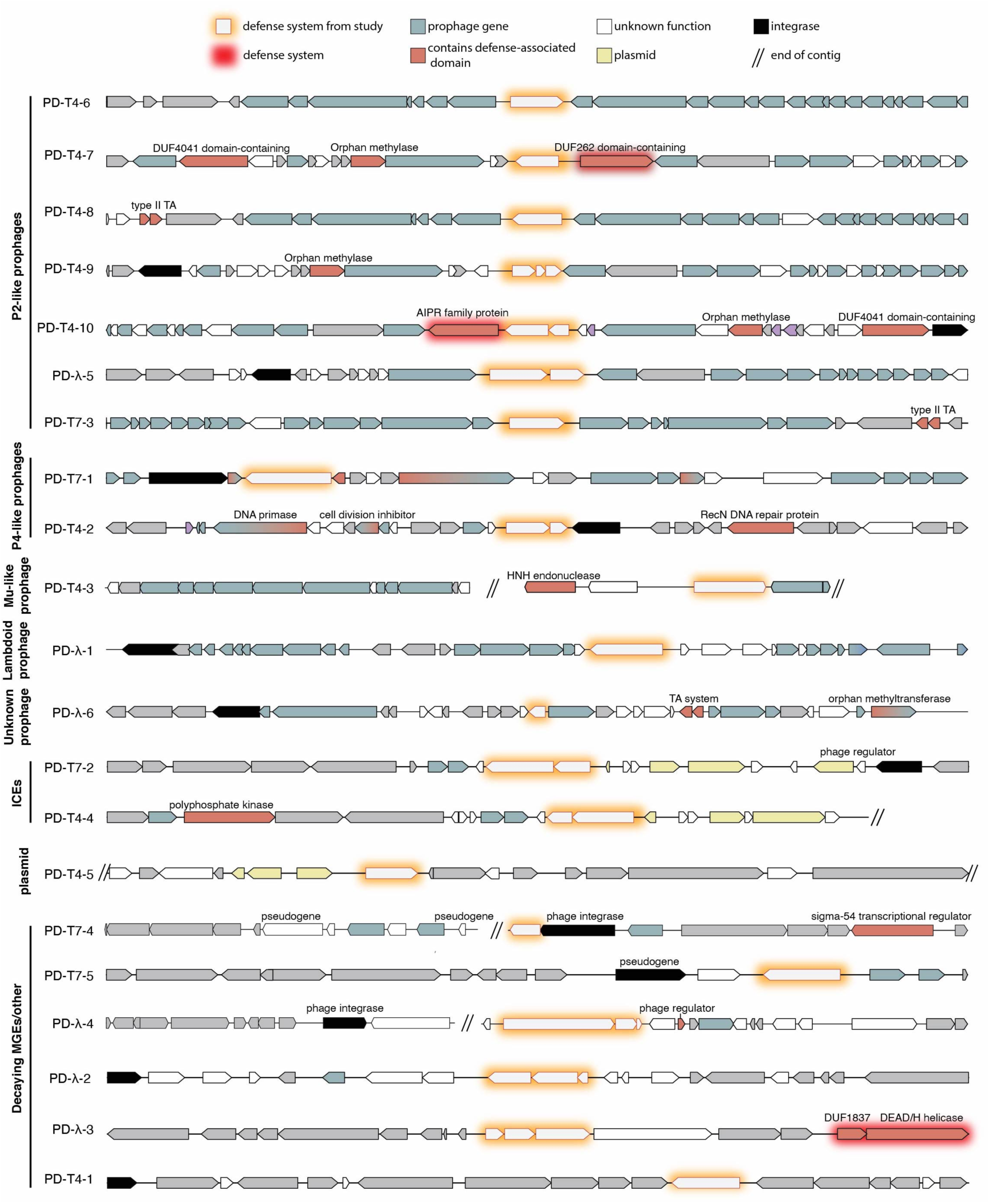
Native genomic locations of defense systems identified. Genome maps of native locations of the defense systems, showing the flanking 10 kb regions, unless interrupted by the end of a contig. Prophage and defense-domain containing genes were called as described in the Methods.

**Figure S5.**
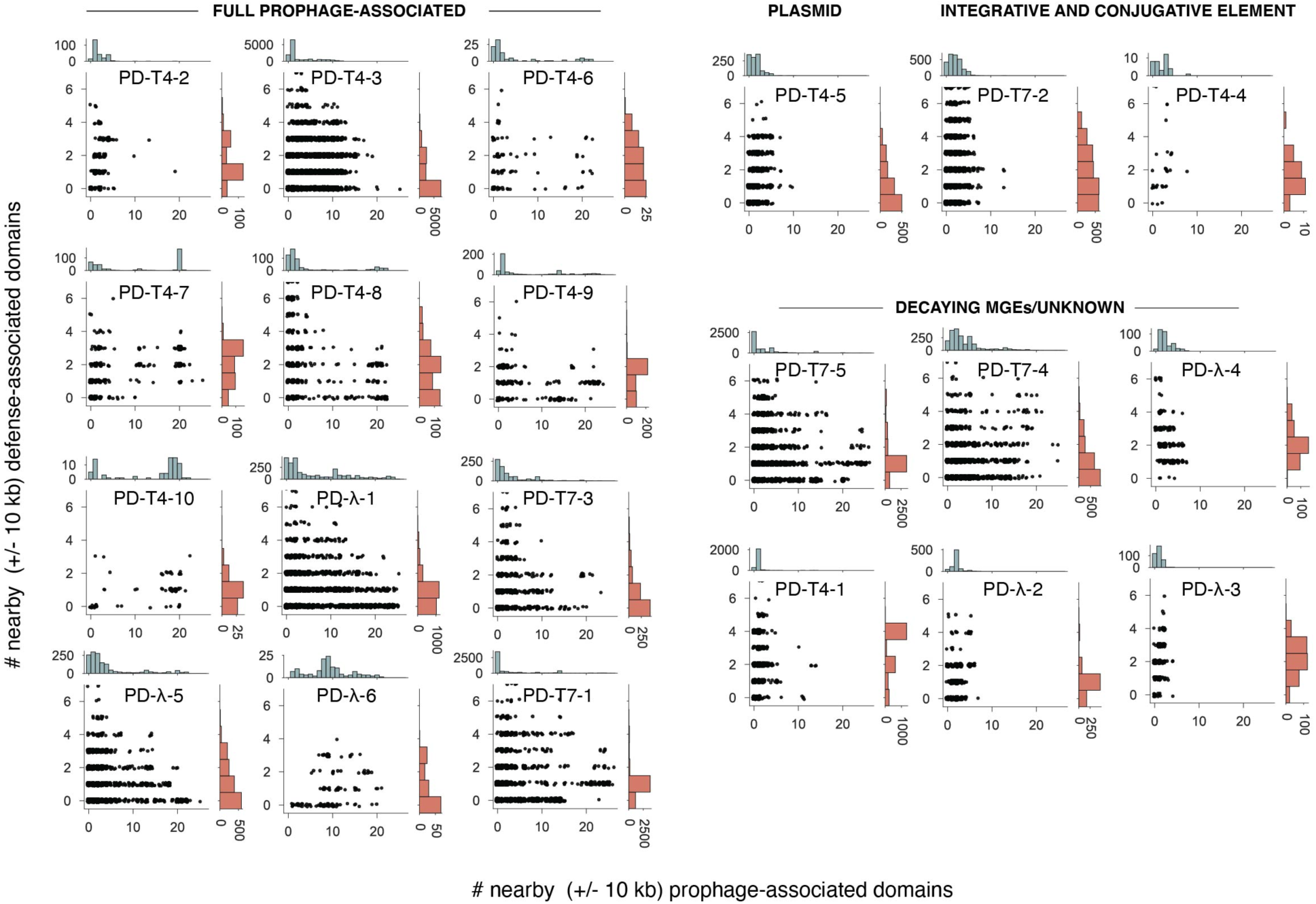
Genome context of defense system homologs and P2 defense-enriched loci. Data as in Fig. 4c, extended to all systems discovered here and sorted by the MGE context in which they were found.

**Figure S6.**
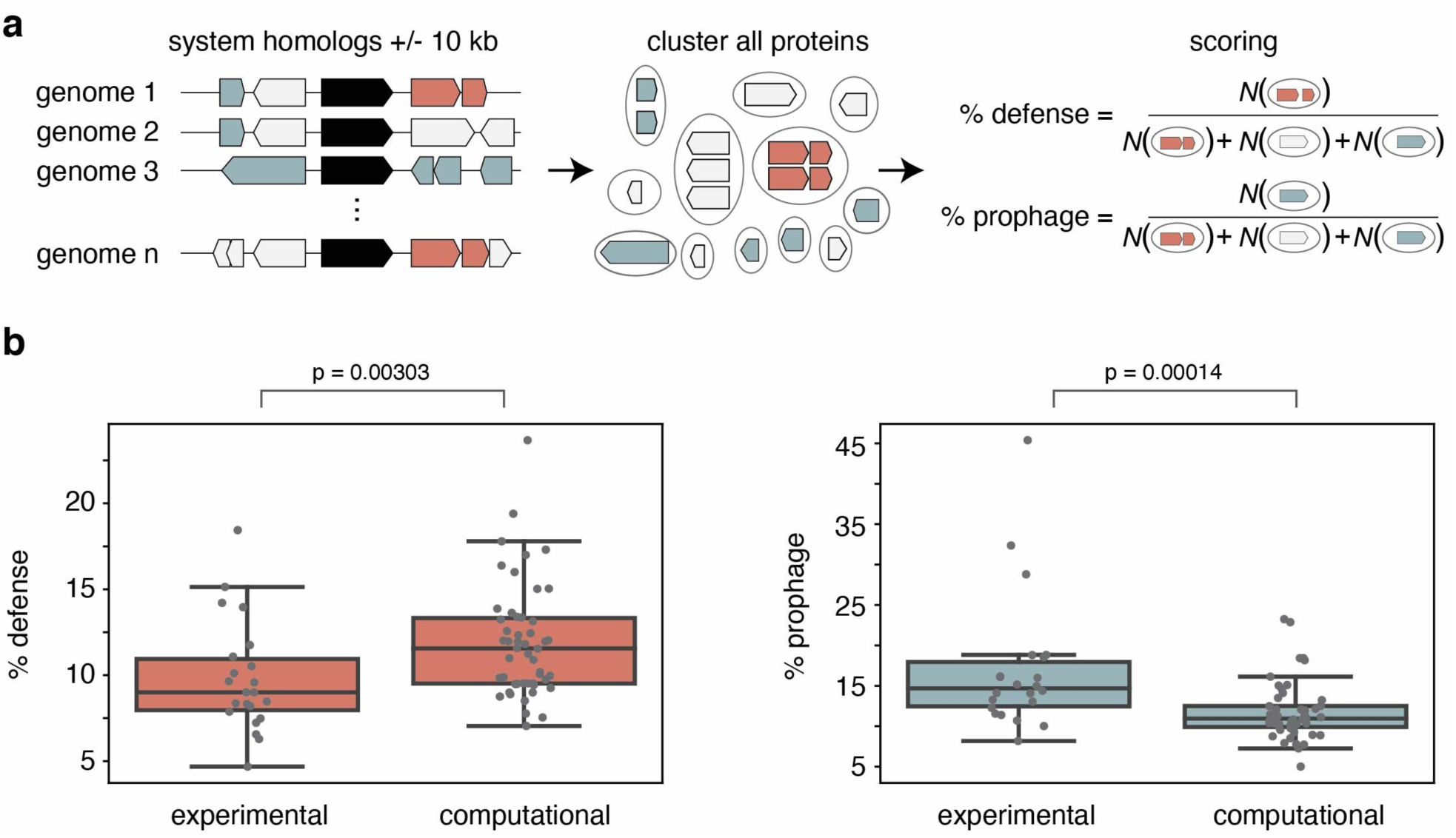
Comparison of defense and prophage enrichments between experimentally and computationally discovered systems. (**a**) Overview of method. Genes +/- 10 kb of homologs of defense systems were predicted as defense- or prophage-associated. To minimize the effects of sequence/genome redundancy, proteins were clustered to 95% identity. % scores were calculated as the number of prophage or defense-associated clusters over total clusters. (**b**) Boxplots of % prophage- and defense-associated genes near the experimentally discovered systems (this study) or computationally predicted and validated systems^12,13^. *p* values indicate significance from Mann-Whitney U test.

